# Deoxynivalenol Promotes Porcine Epidemic Diarrhea Virus Infection and Aggravates Gut Barrier Injury

**DOI:** 10.1101/852608

**Authors:** Dandan Liu, Lei Ge, Qing Wang, Jiarui Su, Xingxiang Chen, Chunfeng Wang, Kehe Huang

## Abstract

Porcine epidemic diarrhea virus (PEDV) is a highly contagious pathogenic virus that causes severe diarrhea and dehydration in pigs of all ages. Deoxynivalenol (DON), the most abundant trichothecene in food and feed, causes vomit and diarrhea in animals and human. However, whether DON exposure could affect PEDV infection remains unknown. Herein, we investigated the impacts of DON on entry and replication of PEDV, morbidity situation of piglets and the mechanisms involved. *In vivo*, twenty-seven piglets infected naturally with PEDV were randomly divided into three groups, receiving the basal diet containing 0, 750 and 1500 μg/kg DON, respectively. We observed significant increases in the diarrhea rates, the villous injury of jejunums and the PEDV proliferation of duodenum, jejunum, ileum and mesenterium of piglets in experimental groups compared with control. Additionally, the autophagosome-like vesicles and the autophagy-related protein expressions were also increased in experimental groups. *In vitro*, we observed that, approximately 2 hrs post-infection, 0.1, 0.5 and 1.0 μM DON promoted PEDV entry (*P* < 0.05) in IPEC-J2s and resulted in tight junction protein occludin internalization. Knockdown of occludin and CRISPR-Cas9-mediated knockout of LC3B indicated a vital role of autophagy-induced occludin internalization in DON-promoted PEDV entry. We also observed that, 24 hrs post-infection, a significant increase in PEDV replication after 0.1, 0.5 and 1.0 μM DON treatment, along with the induction of a complete autophagy. Specifically, deletion of LC3B indicated a crucial role of autophagy in DON-promoted PEDV replication. Pretreatment with SB202190, a p38 signaling inhibitor, abolished the induction of autophagy. Furthermore, downregulation of type I interferon revealed that DON contributed PEDV to escape innate immune. Mechanistically, DON-caused innate immune escape was related to the upregulation of LC3B, which further inhibited STING phosphorylation. Taken together, DON could promote PEDV infection by inducing occludin internalization and innate immune escape via triggering p38-mediated autophagy.

**Author summary:** Porcine epidemic diarrhea (PED), a devastating enteric disease, leads to catastrophic economic loss to the global pig industry. Its primary pathogen is the coronavirus PED virus (PEDV). Growing evidence indicates that pathogen infection is not the only factor of PED outbreaks, other non-infectious factors is also related to this disease. We guessed some ubiquitous substances, such as deoxynivalenol (DON), that lead to pig intestinal epithelial cell stress might encourage the progress and spread of PED. In the present study, the weaning piglets infected naturally with PEDV and the IPEC-J2 cell line were selected as models to explore the effects of DON on PEDV infection, morbidity and gut barrier. Our results showed that DON exposure can promote PEDV infection *in vitro* and *in vivo*, and the underlying mechanism might be related to LC3B-mediated autophagy. Our findings reveal new pathways for developing potential novel antiviral strategies against PEDV infection.

## Introduction

Porcine epidemic diarrhea (PED) is a devastating enteric disease characterized by vomiting, diarrhea and dehydration in pigs of all ages, with up to 90% mortality in suckling piglets, leading to catastrophic economic loss to the global pig industry [1–3]. PED has occurred in China, the United States, Canada, and Vietnam, but its outbreak has a large difference in scale among countries [4–7]. The primary pathogen of PED is PED virus (PEDV), a member of the *Coronaviridae* family. Growing evidence indicates that pathogen infection is not the only cause of PED outbreaks, other non-infectious factors, including stress, management, host intestinal barrier function and immune stimulation, have been suggested to be related to this disease [8–10]. Deoxynivalenol (DON) is a trichothecene mycotoxin that could impair intestinal barrier dysfunction. It is produced by *Fusarium* species and occurs frequently in cereals and animal forages throughout the world, resulting in regular animals and human exposure [11, 12]. Among the farm animals, pigs are the most sensitive species to DON. It has been reported that DON is a major threat to pig health, welfare and performance [13, 14]. However, whether DON contributes to the progress and spread of PED remains unknown.

Autophagy is the major intracellular degradation system that is essential for survival, differentiation and homeostasis [15, 16]. MAP1LC3B/LC3B (microtubule-associated protein 1 light chain 3 β), a marker of autophagic activity, is present during the entirety of this autophagic process and is regulated by lots of signaling [17, 18]. Autophagy principally serves a regulatory mechanism to control the innate immune response against intracellular pathogens [19–21]. On the contrary, in certain viral infection settings, the self-cannibalistic or, paradoxically, even the pro-survival functions of autophagy may be deleterious. Evidence suggest that some viruses, including PEDV, may induce autophagy in order to utilize it for their replication when they infect a target cell [22–24]. However, the impacts of DON on PEDV infection are equivocal, and questions about whether DON induces autophagy in target cells remain unanswered.

In this study, we evaluated the effects of DON on PEDV infection *in vitro* and *in vivo* and found that DON promotes autophagosomes formation, thereby facilitating PEDV replication. Unexpectedly, we also found that DON is required for PEDV entry into the infected cells. Our findings reveal new pathways for developing potential novel antiviral strategies against PEDV infection.

## Results

### Low doses exposure of DON could aggravate intestinal injury and facilitate PEDV infection in weaning piglets

To evaluate the effects of DON exposure on PEDV-infected piglets, we performed animal experiments. Twenty-seven piglets infected naturally with PEDV were randomly divided into three groups: group I received a basal diet, group II received the basal diet containing 750 μg/kg DON; group III received the basal diet containing 1500 μg/kg DON. After 14 days, we observed that the average daily gain (Fig 1A, left) and small intestine weight (Fig 1A, right) of piglets in experiment groups were lower than that in control group (P < 0.05). The diarrhea rates (Fig 1B, left) and diarrhea index (Fig 1B, right) of piglets in experiment groups were increased compared with that in control group (P < 0.05). The pathological results showed that DON exposure aggravates gut barrier injury of PEDV-infected piglets, as demonstrated by the decreases in villus length (Fig 1D, left) and villus/crypt ratio (Fig 1D, right) of jejunums. It’s worth noting that this effect is not caused by DON, as 1500 μg/kg DON exposure alone did not significantly damage the intestinal tract [25]. It also confirmed that 750 and 1500 μg/kg DON were low doses exposure.

**Fig 1.**
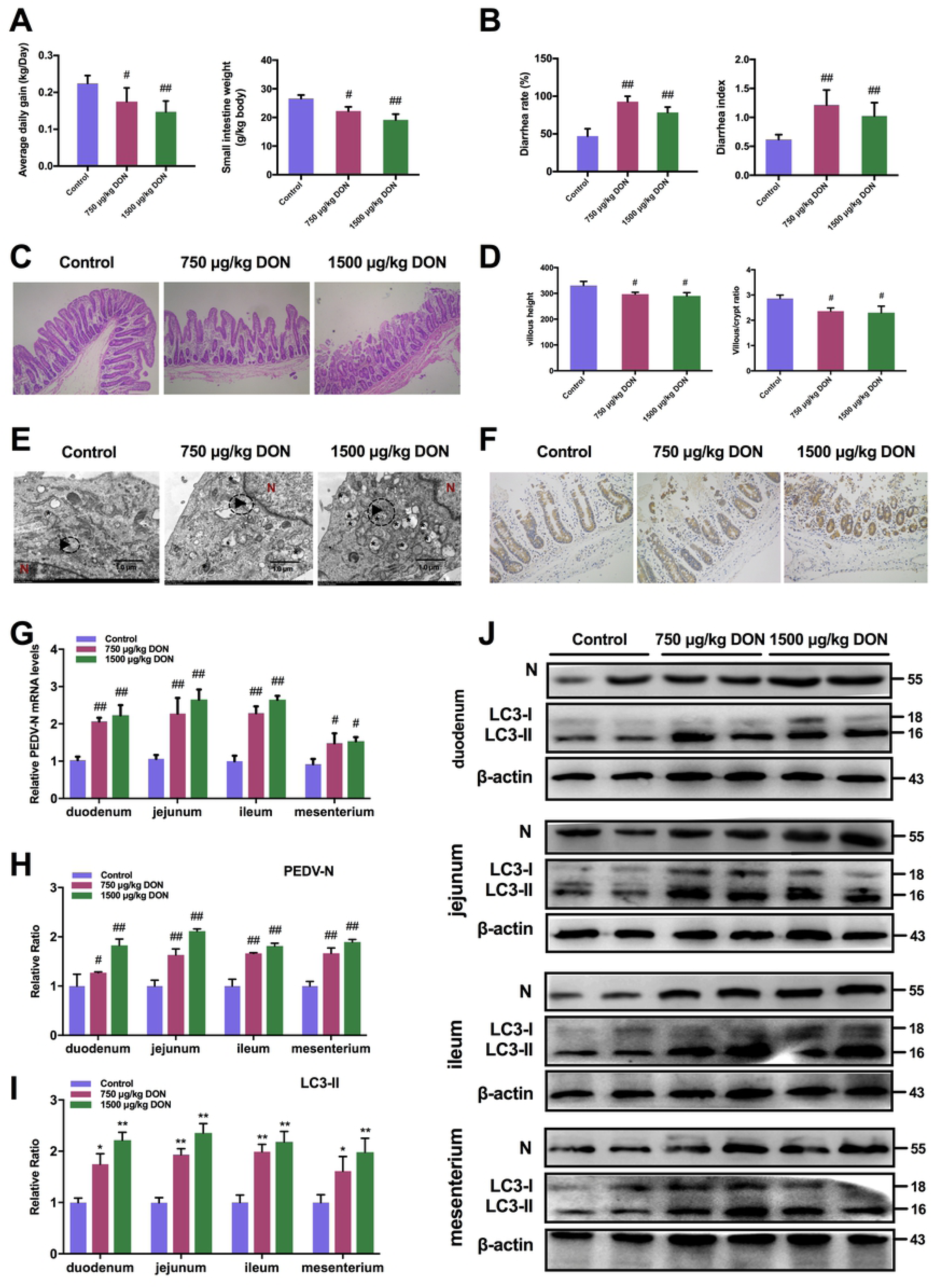
Low concentrations of DON exposure could aggravate intestinal injury and facilitates PEDV infection in weaning piglets. Piglets infected naturally with PEDV were fed with a basal diet containing 0, 750 or 1500 μg/kg DON. (A) Effects of DON on average daily gain and small intestine weight of piglets. (B) Effects of DON on diarrhea rate and diarrhea index of piglets. (C) Histopathological examination. (magnification, × 200) (D) The villus length and villus/crypt ratio of jejunums were quantified. (E) TEM observation. The virus particles (black arrowheads) and the fine ultrastructure of autophagosomes (black asterisk) were observed. The scale bar indicates 1.0 μm. (F) Immunohistochemistry examination. The PEDV-N (brown signals) expression in jejunums of piglets was measured. (magnification, × 200) (G) Effects of DON on the mRNA levels of PEDV-N gene in duodenum, jejunum, ileum and mesenterium. (H, I and J) Effects of DON on the protein levels of PEDV-N and LC3B in duodenum, jejunum, ileum and mesenterium. The data are expressed as mean ± SD (n=3). # *P* < 0.05, ## *P* < 0.01 vs. PEDV. TEM: transmission electron microscopy.

Next, we evaluated that whether low doses exposure of DON could affect PEDV proliferation using transmission electron microscopy (TEM), immunohistochemistry, RT-PCR and immunoblotting. Compared with control group, the virus particles (black arrowheads) observed under TEM were increased in jejunum of piglets in experiment groups (Fig 1E). The immunohistochemistry results showed that the PEDV antigens represented by brown signals in enterocytes of piglets fed with the DON contamination diet were enhanced (Fig 1F). In addition, we found that both the mRNA levels of PEDV-N gene (Fig 1G) and the protein levels of PEDV-N (Fig 1H and 1G) of piglets in experiment groups were increased significantly compared with that in control group. These data suggested that low doses exposure of DON could aggravate intestinal injury of PEDV-infected piglets and facilitate virus proliferation.

### Low doses exposure of DON could trigger autophagy in the intestinal tissues of weaning piglets

To explore whether autophagy can be induced in piglets exposed to DON, the intestinal autophagy levels of piglets in experiment groups were measured by testing the autophagosome-like vesicles formation using TEM and the LC3-II/LC3-I ratio using immunoblotting. As shown in Figure 1E, a larger number of double- or single-membrane vesicles (black asterisk) were observed in jejunum of piglets with the basal diet containing 750 and 1500 μg/kg DON compared with piglets received a basal diet. The LC3-II/LC3-I ratios were also significantly increased in the duodenum, jejunum, ileum and mesenterium of piglets in II and III groups compared with that of piglets in I group (Fig 1I and 1J). These data indicated that DON could trigger autophagy in PEDV-infected piglets, which might be related to DON-promoted PEDV infection as autophagy can facilitate to PEDV proliferation [22].

### Low concentrations of DON could facilitate PEDV entry in IPEC-J2 cells

To eliminate the effects of DON on cytotoxicity, the viability of IPEC-J2 cells treated with different concentrations of DON was analyzed by enzymatic reduction of MTT. As shown in Fig 2A (left), the viability of IPEC-J2 cells was decreased at concentrations of 1.5 to 4.0 μM (P < 0.05). The release of LDH in the supernatant was quantified by detection of LDH enzymatic activity to evaluate the effect of increasing concentrations of DON on the permeabilization of IPEC-J2 cell membrane. Significant increases were observed in the release of LDH after treatment with 1.5 to 4.0 μM DON (Fig 2A, right). Therefore, 0.01, 0.1, 0.5 and 1.0 μM DON were regarded as low concentrations and used in subsequent experiments.

**Fig 2.**
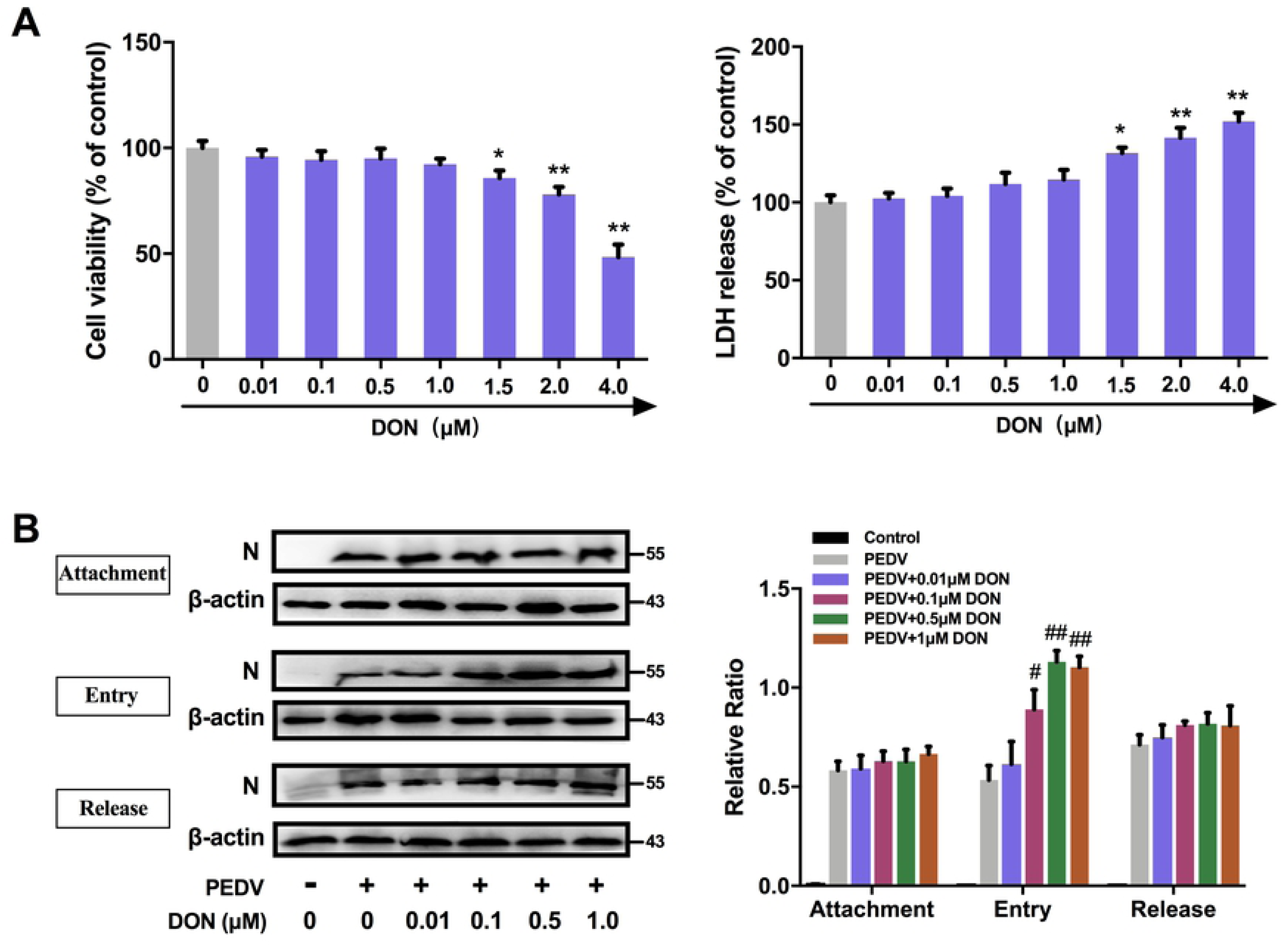
Low concentrations of DON promoted the entry of PEDV in IPEC-J2 cells. (A) Effects of DON on the cell viability and LDH release of IPEC-J2 cells. IPEC-J2 cell monolayers were cultured with various concentrations of DON for 48 h, cell viability and LDH release were assayed as described in Materials and Methods. (B) Effects of low concentrations DON on the PEDV attachment, entry and release. IPEC-J2 cell monolayers were infected with 2 MOI PEDV and further cultured as indicated. Cell lysates were subjected to immunoblotting with antibodies to PEDV nucleocapsid (N) protein or β-actin (loading control). The data are expressed as mean ± SD (n=3). * *P* < 0.05, ** *P* < 0.01 vs. control (mock); # *P* < 0.05, ## *P* < 0.01 vs. PEDV. DON: deoxynivalenol. LDH: lactate dehydrogenase.

To explore whether DON can also affect PEDV infection *in vitro*, we surveyed the relationship between DON exposure and virus attachment, entry and release in IPEC-J2 cells. As determined by immunoblotting, PEDV entry were increased in IPEC-J2 cells exposed to 0.1 - 1.0 μM DON, but there was little change in PEDV attachment and release (Fig 2B). Of note, virus attachment, entry and release revealed no remarkable change following above 1.0 μM DON exposure (Data not shown). These data indicate that low concentrations of DON could facilitate PEDV entry into and release from IPEC-J2 cells.

### Alteration of occludin protein distribution induced by DON contributed to PEDV entry

To explore the mechanism that low concentrations exposure of DON could facilitate PEDV entry, we analyzed the protein levels of the tight junction proteins (ZO-1, occludin and claudin-1) in PEDV-infected IPEC-J2 cells exposed to DON. As determined by immunoblotting, the protein levels of claudin-1 were significantly decreased by 0.5 and 1.0 μM DON and that of ZO-1 had changed little, however, that of occludin were significantly increased by 0.1, 0.5 and 1.0 μM DON in PEDV-infected IPEC-J2 cells (Fig 3A). The cellular expression and distribution of occludin and claudin-1 were measured to further explore the relationship between tight junction proteins and DON-promoted PEDV infection in IPEC-J2 cells. Immunofluorescence analysis (IFA) showed that tight junction formation in mock cells; PEDV infection induced the slight internalization of occludin, not claudin-1, indicating that occludin staining in the junctional area was decreased, and that in cytoplasm was increased; meaning that DON aggravated the internalization of occludin (Fig 3B).

**Fig 3.**
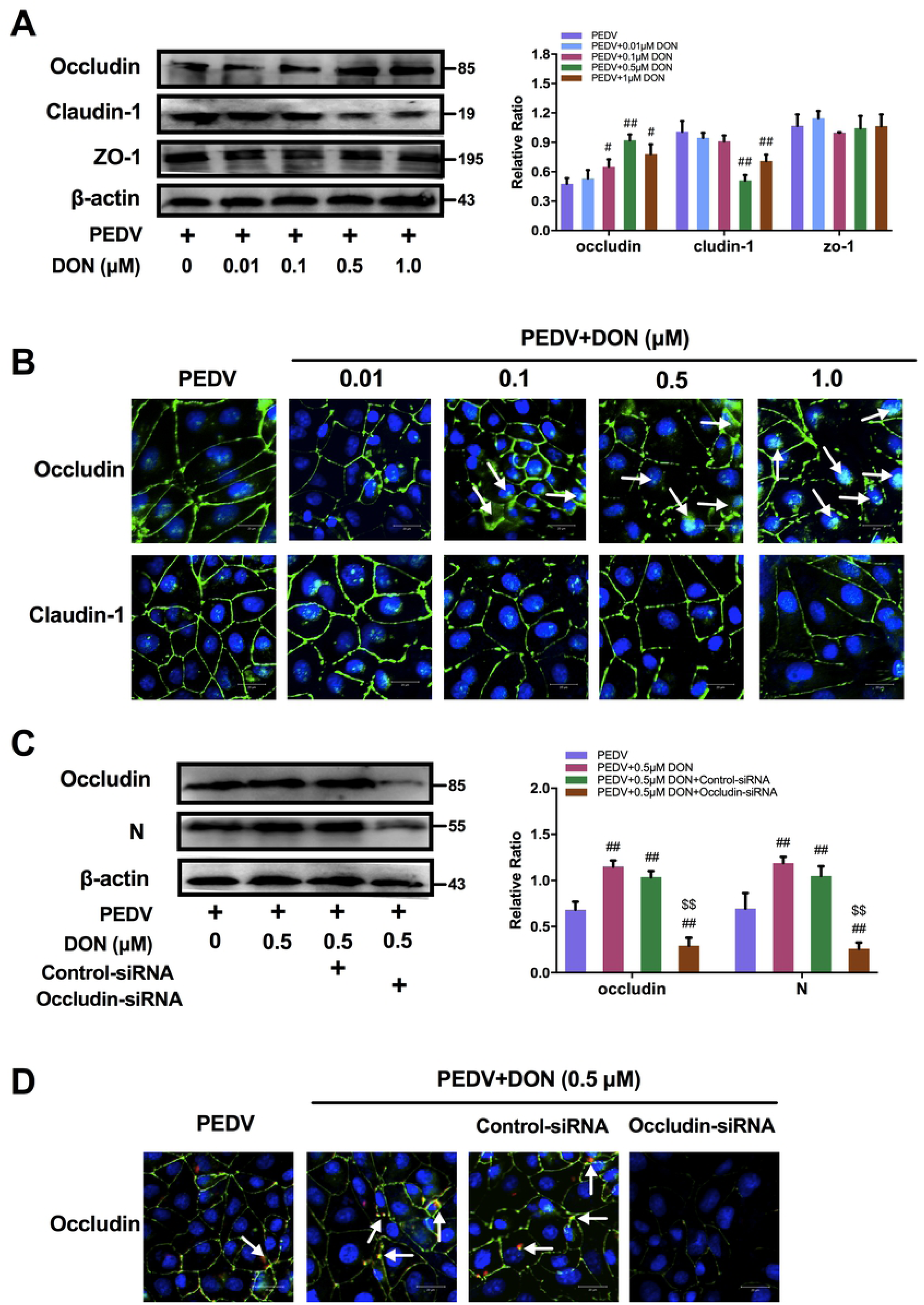
Occludin internalization was required for DON-promoted PEDV entry in IPEC-J2 cells. (A, B) Effects of various concentrations DON on the expression and distribution of tight junction proteins in PEDV-infected IPEC-J2 cells. (C, D) Effects of occludin konckdown on PEDV entry in IPEC-J2 cells exposed to 0.5 μM DON. IPEC-J2 cell monolayers were infected with 2 MOI PEDV and further cultured as indicated. Cell lysates were subjected to immunoblotting (A, C) with antibodies to ZO-1, occludin, claudin-1, PEDV-N protein or β-actin (loading control) and subjected to IFA (B, D) with antibodies to occludin (green), claudin-1 (green) and PEDV-N protein (red). Cell nuclei were stained with DAPI (blue). The scale bar indicates 20 μm. The data are expressed as mean ± SD (n=3). # *P* < 0.05, ## *P* < 0.01 vs. PEDV; $ *P* < 0.05, $$ *P* < 0.01 vs. PEDV+DON. IFA: Immunofluorescence analysis.

Subsequently, small interfering RNA (siRNA) duplexes targeting the occludin gene was used to further determine whether occludin internalization is required for PEDV infection promoted by DON in IPEC-J2 cells. As expected, immunoblotting showed that occludin siRNA-transfected-cells exposed to DON at 0.5 μM were exhibited very low levels of PEDV entry compared with DON+PEDV group (Fig 3C). IFA results validated that occludin knockdown induced a reduction of PEDV-N protein expression (Fig 3D). These data indicate that DON facilitated PEDV entry via altering the cell junctional localization of the occludin.

### CRISPR-Cas9-mediated knockout of the LC3B in IPEC-J2 cells abolished the contribution of DON to occludin-mediated PEDV entry

To explore how DON altered the localization of occludin, we constructed the LC3B-IPEC-J2 cells by CRISPR-Cas9 system to verify whether the LC3B, a hallmark for assessing autophagy, contributed to the occludin localization (Fig 4A). The expression of autophagy-related proteins, occludin and PEDV-N were then measured. The results showed that the expression of LC3-II and the degradation of SQSTM1 were increased significantly in LC3B^+/+^ IPEC-J2 cells treated with DON, which were consistent with the changes of occludin expression. However, the increase in occludin expression and virus proliferation by DON was abolished in LC3B^-/-^ IPEC-J2 cells (Fig 4B). Next, we performed pEGFP transfection assays and observed that the colocalization of LC3B, occludin and virus induced by DON were arrested in LC3B-IPEC-J2 cells (Fig 4C), suggesting that DON induced occludin internalization upon a canonical autophagy.

**Fig 4.**
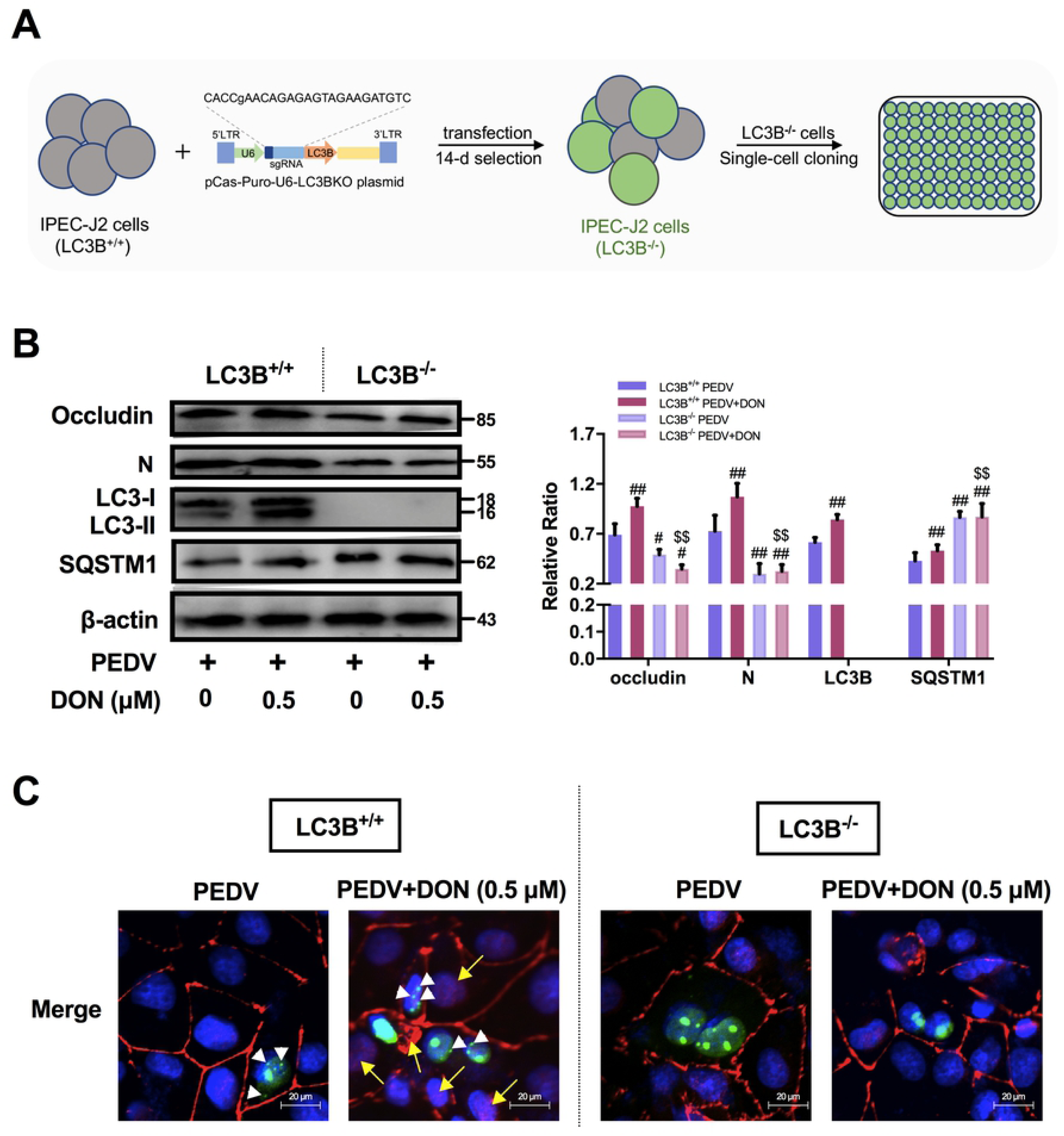
LC3B was required for occludin internalization-induced PEDV entry in IPEC-J2 cells exposed to DON. (A) Generation of LC3B-knockout IPEC-J2 cells. (B, C) Effects of autophagy deficiency on the occludin expression and PEDV entry. IPEC-J2 cell monolayers were infected with 2 MOI PEDV and further cultured as indicated. Cell lysates were subjected to immunoblotting (B) with antibodies to occludin, PEDV-N protein, LC3B or β-actin (loading control). Cell lysates were subjected to IFA (C) with antibody to occludin (red) and plasmid to LC3B (green, white arrowheads). Cell nuclei were stained with DAPI (blue). The scale bar indicates 20 μm. The data are expressed as mean ± SD (n=3). # *P* < 0.05, ## *P* < 0.01 vs. scrambled PEDV; $ *P* < 0.05, $$ *P* < 0.01 vs. scrambled PEDV+DON.

### Low concentrations of DON facilitate PEDV replication in IPEC-J2 cells

To determine the effects of DON on PEDV replication, IPEC-J2 cell monolayers were infected with 1 MOI PEDV for 2 h, and cultured with DON at concentrations between 0.01 and 1 μM for an additional 24 h. The data showed that, compared with the PEDV group, the protein level of PEDV-N (Fig 5A), the viral titer (Fig 5B) and the mRNA levels of PEDV-N and -S genes (Fig 5C) were increased in PEDV-infected cells treated with 0.1, 0.5 or 1 μM DON for 24 h. The maximal effects were observed at 0.5 μM DON. Exposure of cells to DMSO, the solvent control, and 1.5 - 4 μM DON had no effect on PEDV replication (Data not shown). Taken together, these results suggest that low concentrations of DON contributed to PEDV replication in IPEC-J2 cells.

**Fig 5.**
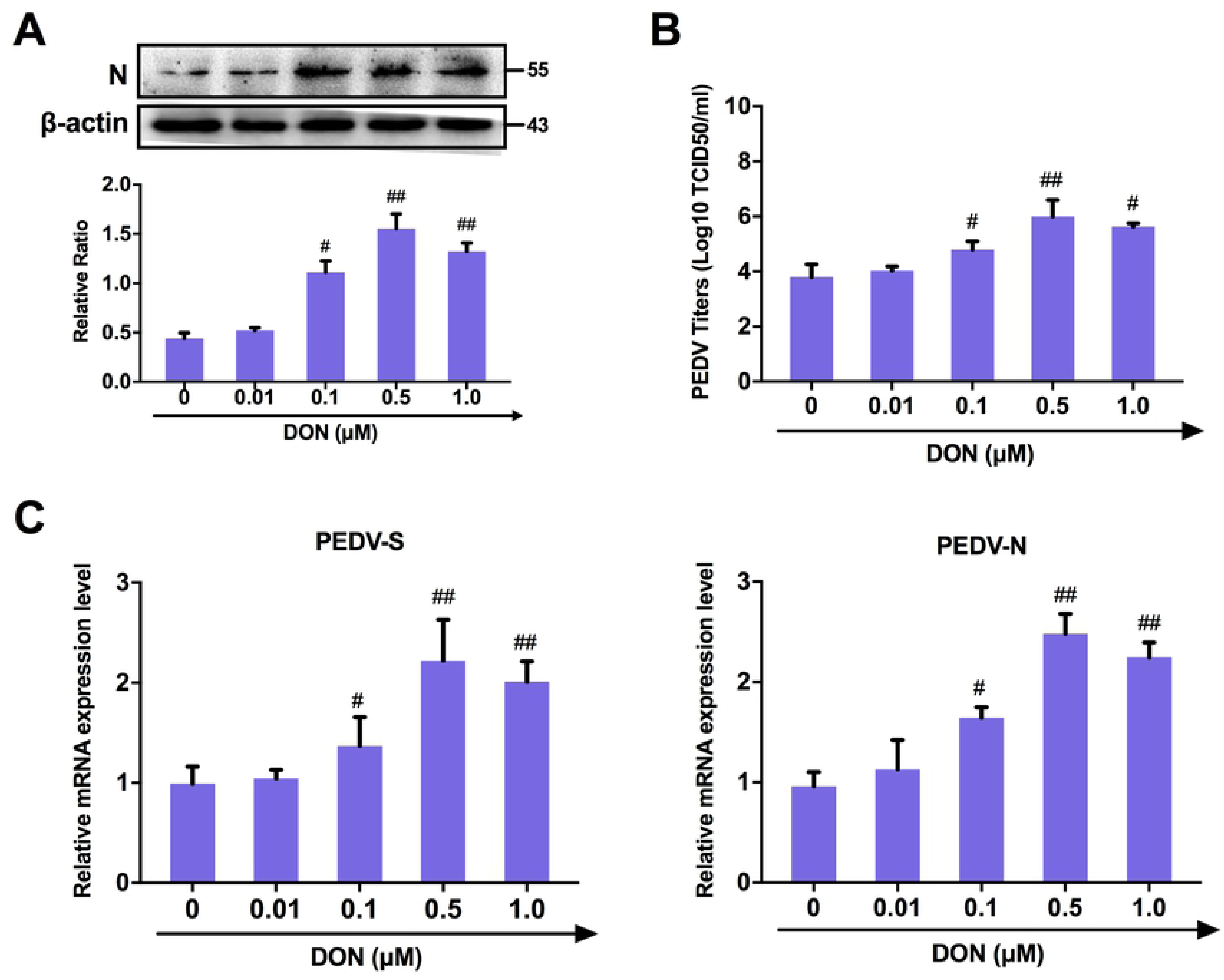
Low concentrations of DON could promote PEDV replication in IPEC-J2 cells. (A) Cell lysates were subjected to immunoblotting with antibodies to PEDV-N protein or β-actin (loading control). (B) Cells were assayed for PEDV viral titers. (C) RT-qPCR were performed to analyze the mRNA levels of PEDV-S and -N genes. The data are expressed as mean ± SD (n=3). # *P* < 0.05, ## *P* < 0.01 vs. PEDV.

### DON triggers a complete autophagic flux in PEDV-infected IPEC-J2 cells

To determine whether autophagy could also play a role in DON-promoted PEDV replication, the level of LC3B was examined and the results showed that DON treatment led to a significant upregulation of LC3-II expression (Fig 6A and 6B). The expression of SQSTM1 was examined to further determine whether a complete autophagic flux was occurred by DON. We found that the protein level of SQSTM1 in PEDV-infected IPEC-J2 cells decreased after DON treatment (Fig 6A and 6B, *P* < 0.05). The maximal effects of DON on the expression of autophagic markers were observed at an DON concentration of 0.5 μM, which is consistent with that in the virus replication result. Moreover, the monomeric red fluorescent protein (mRFP)-Green fluorescent protein (GFP-LC3) tandem reporter construct was used to further measure DON-induced autophagic flux. In the acidic pH of the lysosome, lysosomal hydrolysis can attenuate the green fluorescence of this tandem autophagosome reporter, whereas it has no effect on red fluorescence. Therefore, autophagosomes have both GFP and mRFP signals, whereas autolysosomes have only mRFP signals [26]. As shown in Fig 5D, treatment with CQ, which inhibits the fusion of autophagosomes and lysosomes, resulted in yellow color-labeled autophagosomes, and RFP-LC3-labeled puncta structures were detected in PEDV-infected IPEC-J2 cells expressing the mRFP-GFP-LC3 reporter after incubation with 0.5 μM DON. The similar results could be observed in the immunoblotting experiment (Fig 6A and 6C). These observations indicated that DON induced a complete autophagic flux in PEDV-infected IPEC-J2 cells, which might be responsible for DON-promoted PEDV infection.

**Fig 6.**
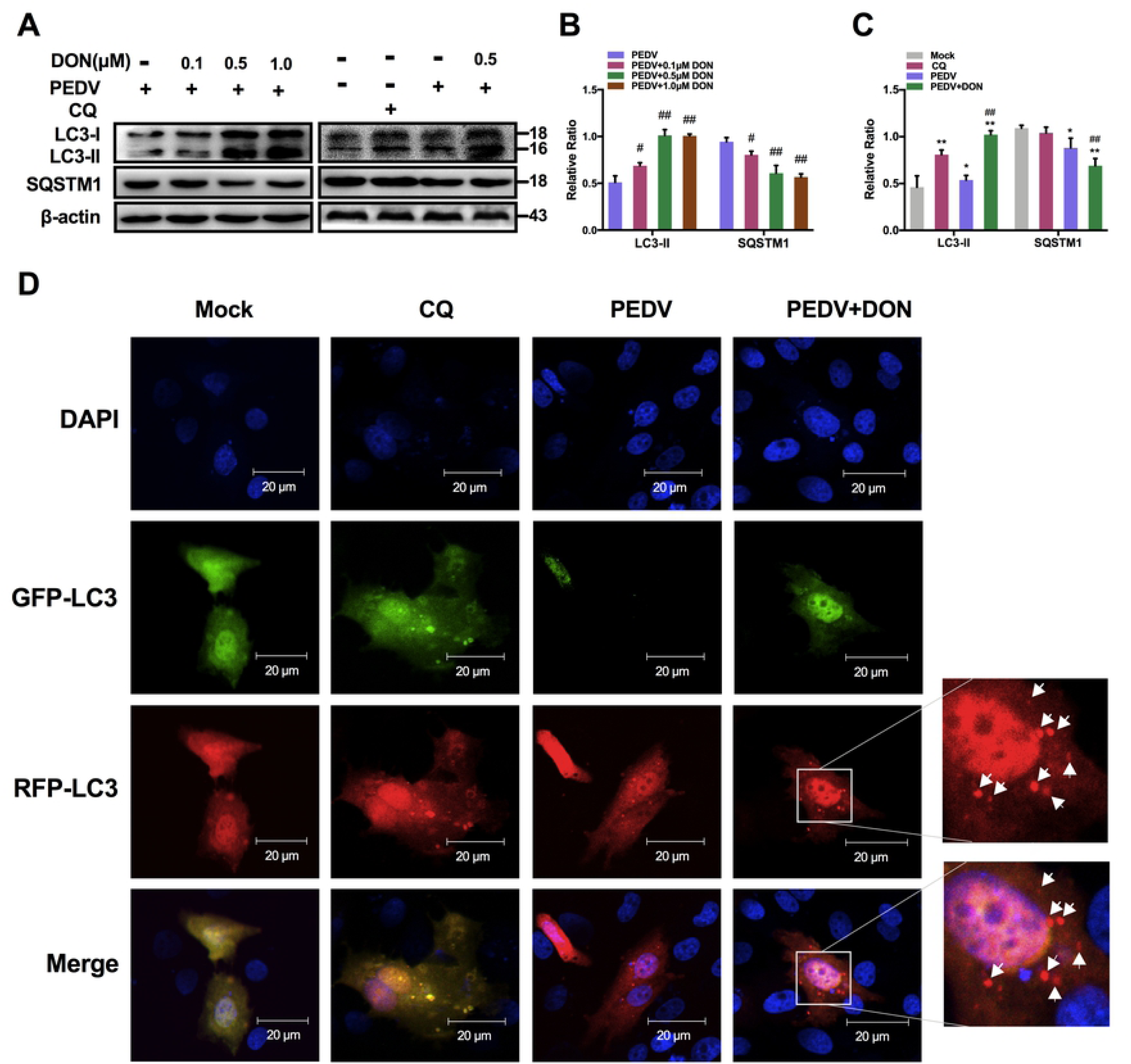
Low concentrations of DON could promote autophagosomes formation in PEDV-infected IPEC-J2 cells and piglets. Cell monolayers were infected with 1 MOI PEDV for 2 h, and cultured with or without DON for another 24 h. (A, B and C) Cell lysates were subjected to immunoblotting with antibodies to autophagy-related proteins (LC3B and SQSTM1) or β-actin (loading control). (D) Cell lysates were subjected to IFA with plasmid to autophagosomes (green spots) and autophagolysosomes (red spots, white arrowheads). Cell nuclei were stained with DAPI (blue). The scale bar indicates 20 μm. The data are expressed as mean ± SD (n=3). * *P* < 0.05, ** *P* < 0.01 vs. control (mock); # *P* < 0.05, ## *P* < 0.01 vs. PEDV. CQ: chloroquine.

### CRISPR-Cas9-mediated knockout of LC3B in IPEC-J2 cells suppressed the promotion of DON to PEDV replication

To further confirm the role of autophagy in DON-promoted PEDV replication, we compared the viral yield in LC3B^+/+^ and LC3B^-/-^ IPEC-J2 cells exposed to DON. A significant decrease was observed in the protein expression of LC3B in LC3B^-/-^ IPEC-J2 cells compared with that in LC3B^-/-^ cells. Moreover, PEDV viral yield exhibited the same decrease, as demonstrated by the down-regulation of PEDV-N protein level (Fig 7B), PEDV viral titers (Fig 7C), and PEDV-N / -S mRNA levels (Fig 7D), indicating that the decreased viral yield was due to the inhibition of autophagy. Collectively, these results suggest that the LC3B-medicated autophagy machinery was required for DON-promoted PEDV replication in IPEC-J2 cells.

**Fig 7.**
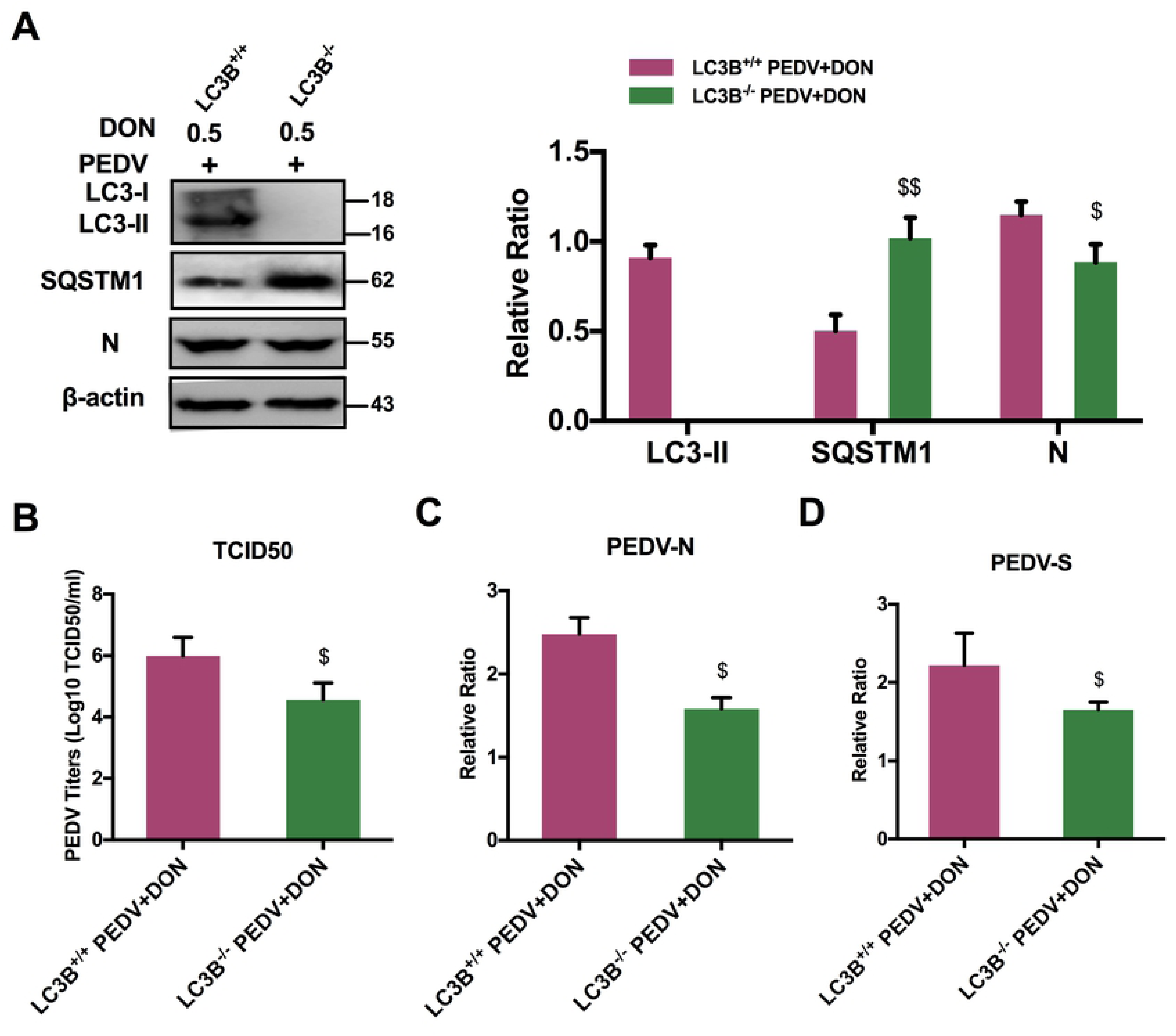
Inhibition of autophagy could decrease DON-promoted viral yield in IPEC-J2 cells. Cell monolayers were infected with 1 MOI PEDV for 2 h, and cultured with or without DON or shLC3B for another 24 h. (A) Cell lysates were subjected to immunoblotting with antibodies to LC3B, SQSTM1, PEDV-N or β-actin (loading control). (B) Cells were assayed for PEDV viral titers. RT-qPCR were performed to analyze the mRNA levels of PEDV-N (C) and -S (D) genes. The data are expressed as mean ± SD (n=3). * *P* < 0.05, ** *P* < 0.01 vs. control (mock); # *P* < 0.05, ## *P* < 0.01 vs. PEDV; $ *P* < 0.05, $$ *P* < 0.01 vs. PEDV+DON.

### Activation of p38/MTORC1 signaling pathway was required for the upregulation of LC3B by DON in PEDV-infected IPEC-J2 cells

To explore how DON induced autophagy, JAKs, PI3K and MAPKs signaling related-proteins were detected. The immunoblotting results revealed that there was no significance in JAK1, PI3K, JNK/p-JNK and ERK/p-ERK proteins expression after DON treatment. But, a significant increase in p-p38 was observed (Fig 8A). In addition, p-MTORC1 was significantly downregulated by DON. So, we supposed the activation of p-p38 might induce autophagy. To determine our hypothesis, the inhibitor of p-p38, SB202190, was supplied. The data showed that SB202190 inhibited the LC3II activation and SQSTM1 degradation induced by DON, upregulated the p-MTORC1 expression (Fig 8B) and blocked the formation of autophagosomes (Fig 8C). Therefore, DON induced autophagy via upregulating the p38/MTORC1 signaling pathway in PEDV-infected IPEC-J2 cells.

**Fig 8.**
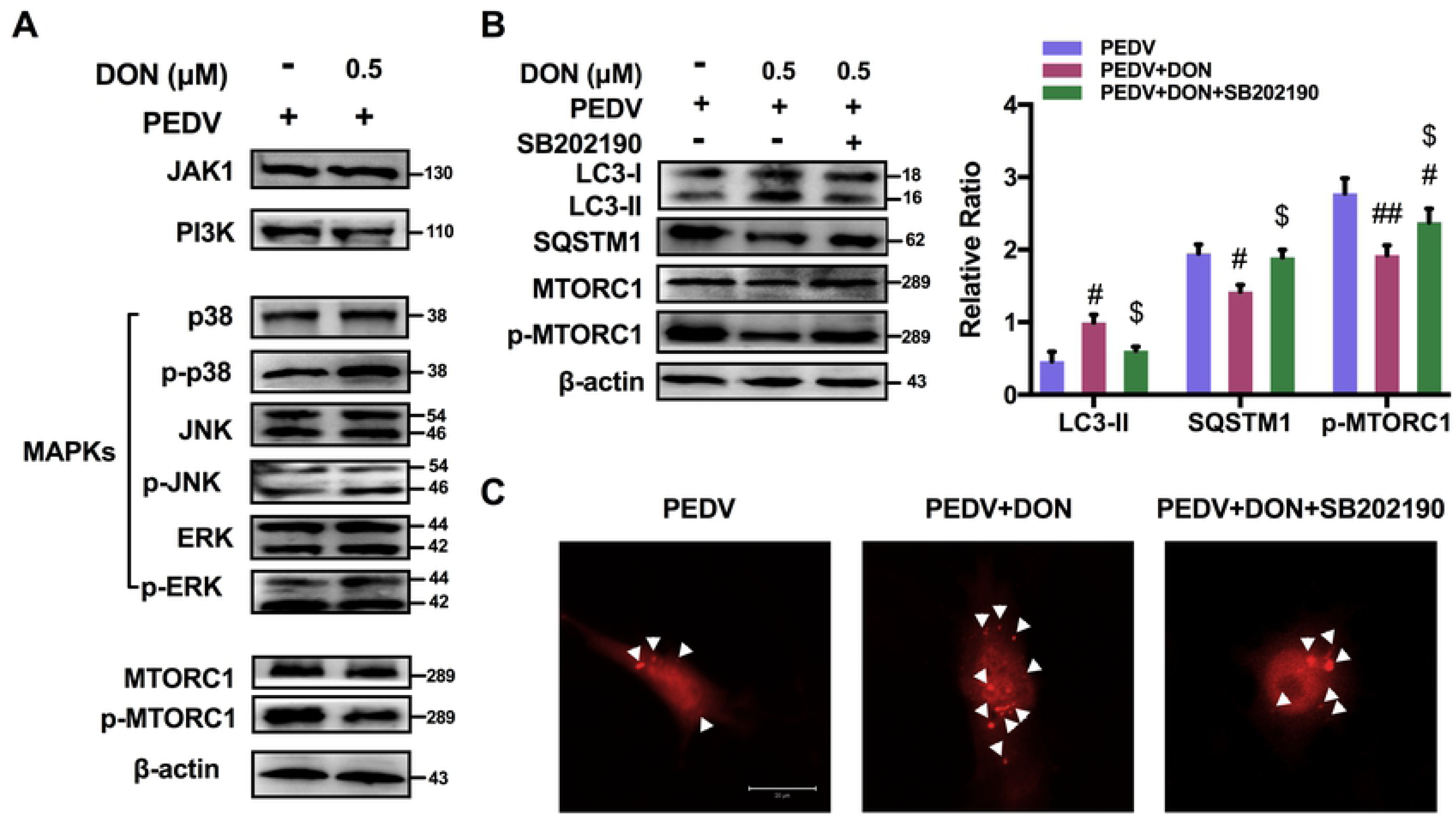
Activation of p38/MTORC1 signaling pathway was required for the activation of LC3B-mediated autophagy by DON in PEDV-infected IPEC-J2 cells. Cell monolayers were infected with 1 MOI PEDV for 2 h, and cultured with or without DON for another 24 h. (A) Cell lysates were subjected to immunoblotting with antibodies to JAK1, PI3K, MAPKs, p-MTORC1/MTORC1 or β-actin (loading control). (B) Cell lysates were subjected to immunoblotting with antibodies to LC3B, SQSTM1, p-MTORC1/MTORC1 or β-actin (loading control) at the present of p-p38 inhibitor, SB202190. (C) Cell lysates were subjected to confocal with plasmid to autophagosomes (red spots, white arrowheads). The scale bar indicates 20 μm. The data are expressed as mean ± SD (n=3). # *P* < 0.05, ## *P* < 0.01 vs. PEDV; $ *P* < 0.05, $$ *P* < 0.01 vs. PEDV+DON.

### Low concentrations of DON facilitate PEDV to escape innate immune by activating autophagy in IPEC-J2 cells

To explore how autophagy affected virus replication, the effects of DON on interferon (IFN-α, IFN-β, IFN-γ, and IFN-λ) expression in PEDV-infected IPEC-J2 cells were measured as autophagy is an important component of both innate and acquired immunity to pathogens [27]. Following poly (I:C) transfection, 0.1, 0.5 and 1 μM DON treatment specifically inhibited the expression of IFN-α and IFN-β compared with PEDV-infected cells (Fig 9A and 9B). However, there was no significance in the mRNA level of IFN-γ and IFN-λ (Fig 9C and 9D) after DON treatment. The shLC3B was used to confirm whether autophagy played a role in downregulation of type I interfere. The results showed that DON-downregulated type I interfere could be blocked by shLC3B (Fig 9E and 9F, *P* < 0.05). These data indicated that autophagy played a key role in the suppression of antiviral innate immune by DON.

**Fig 9.**
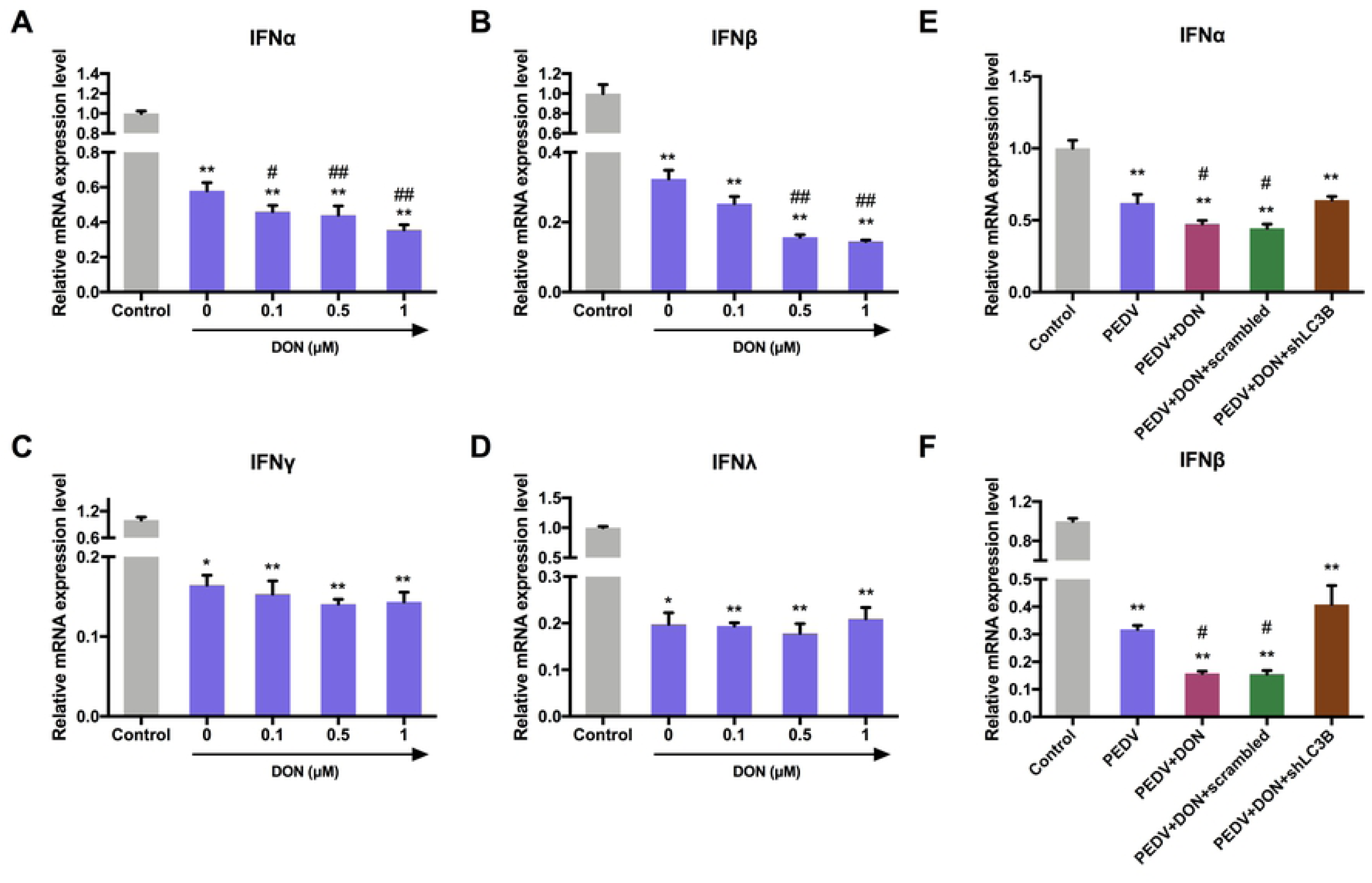
Low concentrations of DON could mitigate the antiviral innate immune response by activating autophagy in PEDV-infected IPEC-J2 cells. Cell monolayers were infected with 1 MOI PEDV for 2 h, and cultured with or without DON for another 24 h. RT-qPCR were performed to analyze the mRNA levels of IFN-α (A), IFN-β (B), IFN-γ (C) and IFN-λ (D). RT-qPCR were performed to analyze the mRNA levels of IFN-α (E) and IFN-β (F) after treating with autophagy inhibitor, 3-MA. The data are expressed as mean ± SD (n=3). * *P* < 0.05, ** *P* < 0.01 vs. control (mock); # *P* < 0.05, ## *P* < 0.01 vs. PEDV.

### Autophagy-mediated STING pathway was required for PEDV escape innate immune in IPEC-J2 cells treated with DON

To explore how autophagy participated in the regulation of innate immune, STING signaling was detected in PEDV-infected IPEC-J2 cells treated with DON. The immunoblotting results revealed that DON treatment inhibited the phosphorylation of STING (Fig 10A, *P* < 0.01), which was consistent with the changes in expression of IFN-α and IFN-β. So, we supposed that autophagy downregulated the expression of type I interfere via inhibiting STING signaling pathway. The scrambled and shLC3B plasmids were constructed and used to determine our hypothesis. The data showed that shLC3B significantly blocked the inhibitory effects of DON on STING phosphorylation (Fig 10B). In addition, the inhibition of type I interfere expression were also significantly blocked (Fig 10C and 10D), suggesting that STING signaling was required to control the replication promotion of PEDV by DON and its antiviral activities were mediated by autophagy.

**Fig 10.**
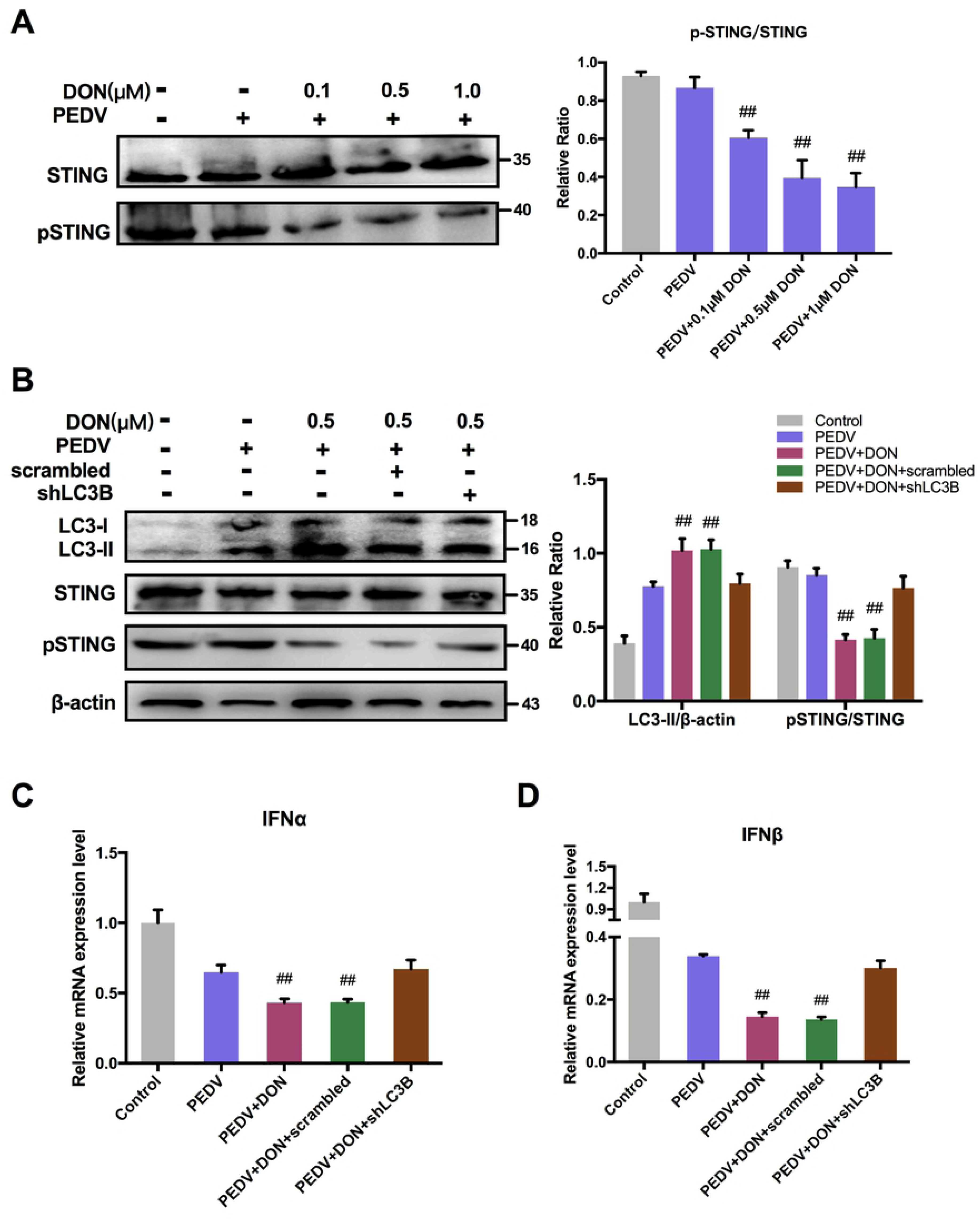
Activation of autophagy by DON suppressed the antiviral innate immune response via inhibiting STING signaling phosphorylation in PEDV-infected IPEC-J2 cells. Cell monolayers were infected with 1 MOI PEDV for 2 h, and cultured with or without DON for another 24 h. (A) Cell lysates were subjected to immunoblotting with antibodies to p-STING and STING. (B) Cell lysates were subjected to immunoblotting with antibodies to LC3B, p-STING/STING or β-actin (loading control) at the present of scrambled or LC3B. RT-qPCR were performed to analyze the mRNA levels of IFN-α (C) and IFN-β (D) at the present of scrambled or LC3B. The data are expressed as mean ± SD (n=3). # *P* < 0.05, ## *P* < 0.01 vs. PEDV.

**Fig 11.**
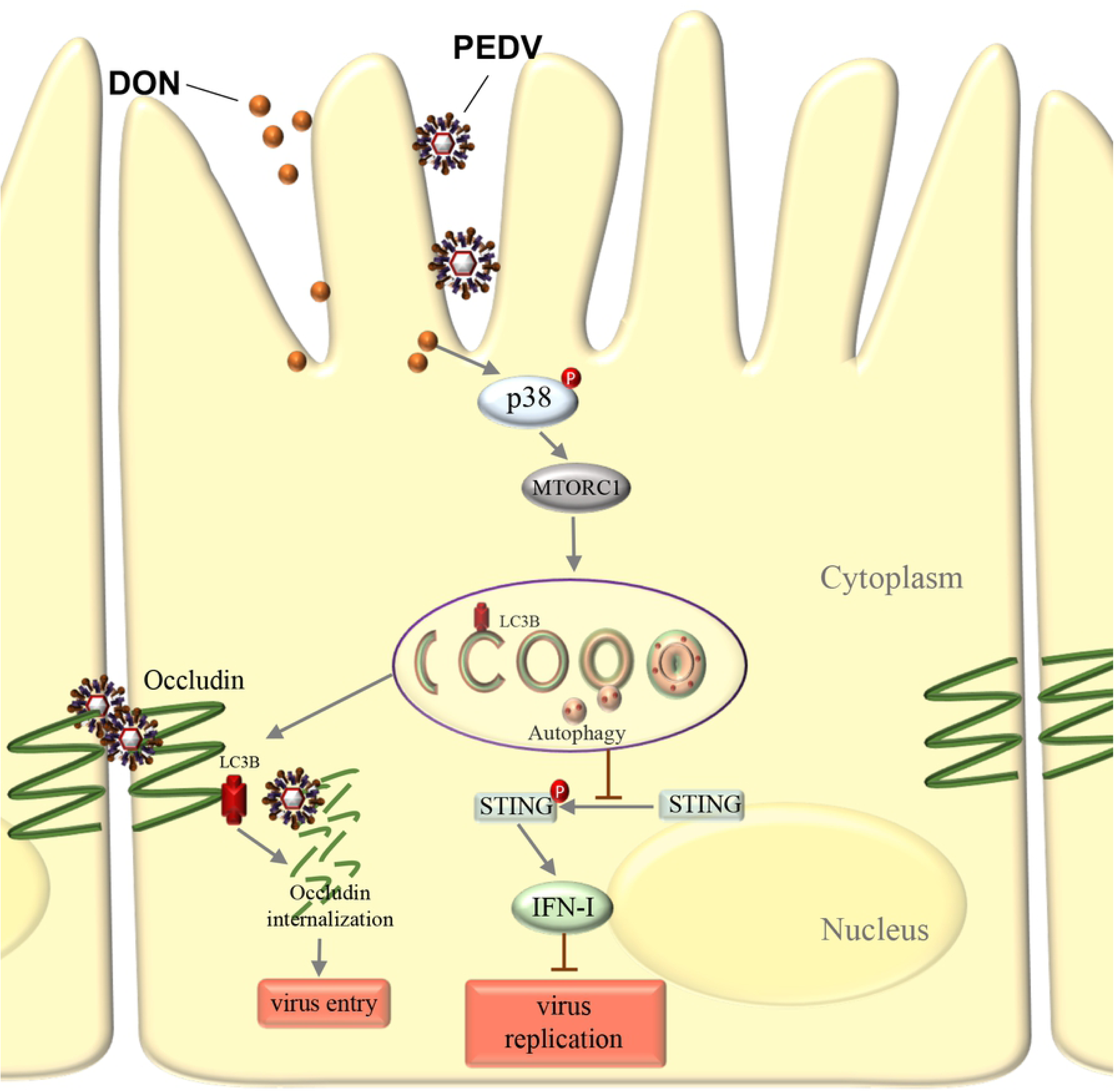
Schematic depicting role of LC3B during PEDV infection. DON exposure activates p38 signaling and triggers a complete autophagy in PEDV-infected IPEC-J2 cells. Approximately 2 hrs post-infection, LC3B induces occludin internalization to promote PEDV entry through its role as a positive regulator of autophagy. Later in infection (24 hpi.), the up-regulation of LC3B by DON contributes PEDV to escape innate immune via inhibiting the STING signaling phosphorylation, leading to production of large amounts of virus.

## Discussion

The contamination of foods and feeds with DON is a significantly serious problem worldwide [28–30]. Pigs are considered to be one of the most sensitive species. They are frequently exposed to DON owing to grains account for a large proportion in their feedstuffs. PED outbreaks caused by PEDV is now distributed all over the world [31]. Thus, the co-existence of DON and PEDV occurs frequently in global pig farms. However, whether DON exposure may increase the susceptibility to PEDV remains unknown. In this study, we provide the first strong evidence that DON exposure can promote PEDV infection *in vitro* and *in vivo*, and the underlying mechanism might be related to LC3B-mediated autophagy.

Intestinal mucosal, the first barrier to food contaminants, chemicals, and pathogens, plays an important role in regulating the immune response to these stressors [32, 33]. Since the ability of DON to efficiently cross biological barriers, fast dividing cells such as intestinal epithelial cells will be more susceptible to the detrimental effects of DON [11, 34]. And PEDV mainly infects pig small intestinal epithelial cells. We then hypothesized that some substances, such as DON, that lead to pig intestinal epithelial cell stress might encourage the progress and spread of PED. Therefore, the intestinal porcine epithelial cell line IPEC-J2 was used as an *in vitro* model of swine small intestine epithelium. Experiments *in vivo* were performed on twenty-seven weaning piglets received the basal diet containing DON. In the present study, we found, for the first time, that low concentrations of DON could facilitate PEDV infection *in vivo* and *in vitro*.

What drives DON to promote virus infection? PEDV crosses the porcine intestinal mucosa to cause intestinal infection, and then results in an acute viral enteric disease, which means that PEDV must gain access to the tight junctions. As noted earlier, the alteration of tight junction proteins distribution might participate in virus entry, for example, occludin internalization contributes to PEDV entry [35]. In the experiments reported here, we demonstrated that DON could aggravate occludin internalization in PEDV-infected cells. When the occludin gene was silenced by siRNA, the promotion of DON to PEDV entry in IPEC-J2 cells was disappeared simultaneously, indicating that occludin internalization contributes to the DON-induced PEDV entry. And what was responsible for occludin internalization was further confirmed.

Autophagy is a selective degradation process of various subcellular structures, including protein aggregates. Increasing evidence indicated that autophagy is related to cell membrane integrity and membrane proteins distribution [36]. It can serve dual roles in virus infection with either pro- or anti-viral functions depending on the virus and the stage of the viral replication cycle [37]. It not only is required for an antiviral response against some virus infection [38], but also take an active part in the viral life cycle by, eg, facilitating its entry into and release from cells [39]. In PEDV-infected cells, autophagy is often hijacked by viruses and manipulated to their own advantage [22]. Our data in this study have showed that the autophagy levels were significantly increased by DON *in vitro* and *in vivo*, which are consistent with the viral infection levels. CRISPR-Cas9-mediated knockout of the LC3B blocked the promotion of DON to PEDV viral yield. Therefore, we concluded that autophagy is required for DON-promoted PEDV infection, including virus entry and replication. In addition, previous studies indicated that induction of autophagy is associated with enhanced JAK1 [40], PI3K and MAPKs signalings [41–43]. Our study confirmed that MAPK p38 signaling was enhanced in PEDV-infected IPEC-J2 cells treated with DON, accompanied by a decrease in mTORC1 levels.

But, how did DON-activated autophagy promote PEDV replication? The primary role of autophagy in innate immune is regulating the IFN-I expression [44, 45]. It is well known that IFN-I is an important antiviral defense cytokine in innate immunity. Upregulation of IFN-I can inhibit viral proliferation, whereas downregulation of it contributes to virus infection [46]. PEDV infection is one of the main mechanisms for inhibiting IFN-I signaling during continuous infection [47]. However, questions that whether DON exposure could facilitate down-regulation of IFN-I in PEDV-infected cells remain unanswered. We investigated the effects of DON to IFN-I expression and confirmed that low concentrations of DON facilitated PEDV to escape innate immune by activating autophagy in IPEC-J2 cells. Stimulator of interferon genes (STING, TMEM173, MITA) is a critical component of the cellular innate immune response to pathogenic cytoplasmic DNA and expressed predominantly in the endoplasmic reticulum (ER) [48]. It can be activated by the enzyme cGAMP synthase (cGAS) and then activates interferon regulatory factors (IRFs) and NF-κB, which leads to the induction of type I interferon and other immune response genes. We then investigated the contribution of STING to autophagy-inhibited IFN expression for that attenuation of the STING signaling can occur through autophagy [49] and confirmed that autophagy-mediated STING pathway played a crucial role in the innate immune escape of PEDV in IPEC-J2 cells treated with DON.

Current PEDV pathogenesis target primarily virus infection, little attention focus on non-infectious factors. This study provides evidence for the first time that low concentrations of DON can promote PEDV infection *in vitro* and *in vivo*, documenting that autophagy activated by DON modulates the promotion. The present study also provides new insight into developing potential novel antiviral strategies against PEDV infection.

## Materials and methods

### Animal experiments

All experiments were conducted according to the standards of the European Guidelines for Animal Welfare and were approved by the Committee for the Care and Use of Experimental Animals of the Nanjing Agricultural University (Animal Ethics Number: SYXK (Su) 2011-0036). Animal experiments were carried out at a 1400-weaning piglets farm. Eighty weaning piglets (age 3 weeks) were selected. These piglets were positive for PEDV naturally and negative for transmissible gastroenteritis virus and porcine rotavirus as determined by fecal and blood diagnostics.

Twenty-seven out of eighty piglets (BW= 5.5 ± 0.5 kg) were selected and randomly divided into three groups, with 3 replicates per group and 3 piglets per replicate. Piglets were assigned to 3 groups (I: control, received a basal diet, II: received the basal diet containing 750 μg/kg DON, and III: received the basal diet containing 1500 μg/kg DON), for 14 days. All piglets were housed in the same facility but different rooms under biosafety conditions and allowed free access to water and feed during the experiment. Body weight and feed intake were recorded to determine the growth performance of piglets. On dpi 14, the piglets were euthanized and tissue samples of duodenum, jejunum, ileum and mesenterium were collected.

### Diarrhea rate and diarrhea index evaluation

Diarrhea in each pig was recorded and scored daily according to the state of feces. Piglets with dry and cylindrical feces are scored 0 point. Piglets with soft and tangible feces are scored 1 point. Piglets with sticky and semi-solid feces are scored 2 points. Piglets with liquid and unformed feces are scored 3 points. Diarrhea rate = [number of diarrhea piglets per replicate / (number of piglets per replicate * days)] * 100%. Diarrhea index = diarrhea scores sum of piglets of per replicate / (number of piglets per replicate * days).

### Reagents and antibodies

Deoxynivalenol (DON, purity≥98%, for experiments *in vitro*), 3-methlyadenine (3-MA), chloroquine (CQ) and rabbit anti-LC3B antibody were purchased from Sigma-Aldrich (St. Louis, USA). Deoxynivalenol (DON, purity≥98%, for experiments *in vivo*) was purchased from Pribolab (Immunos, Singapore). Rabbit anti-SQSTM1, anti-MAPKs, anti-JAK1, anti-pSTING/STING, anti-PI3K, anti-β-actin antibodies and horseradish peroxidase (HRP)-conjugated goat anti-rabbit secondary antibody were purchased from Cell Signaling Technology (Boston, USA). Rabbit anti-claudin1, anti-occludin, anti-ZO-1 and anti-p-MTORC1/MTORC1 antibodies were purchased from Abcam (Cambridge, UK). Porcine epidemic diarrhea virus (PEDV) strain CV777 was obtained from Jiangsu Academy of Agricultural Sciences (Nanjing, China). Rabbit anti-PEDV-N antibody was prepared by our lab. Poly (I:C) (LMW) / LyoVec^TM^ was purchased from InvivoGen (San Diego, USA). SB203580 was purchased from MCE (New Jersey, USA).

### Histological analysis

Jejunum tissues samples were fixed in 4% paraformaldehyde, embedded in paraffin and sectioned at a thickness of 4 μm. For histopathological examination, tissue slices were stained with hematoxylin and eosin, and observed under the microscope. For immunohistochemistry examination, tissue slices were incubated with antibodies against PEDV N protein, followed by incubation with the second antibody and streptavidin-peroxidase complex. The peroxidase conjugates were visualized using DAB solution.

### Quantitative Real-Time PCR (RT-qPCR) Analysis

RT-qPCR was performed using the StepOnePlus Real-Time PCR System (Applied Biosciences) as previously described [50]. Total RNA was isolated from the cells using an RNA Extraction Kit (Takara, Japan) and then reverse-transcribed to cDNA using the PrimeScript RT Master Mix Kit (Takara, Japan). All primers are listed in Table 1. The relative expression was determined using the Δcycle threshold (ΔC_t_) method with GAPDH serving as a reference gene.

**Table 1.**
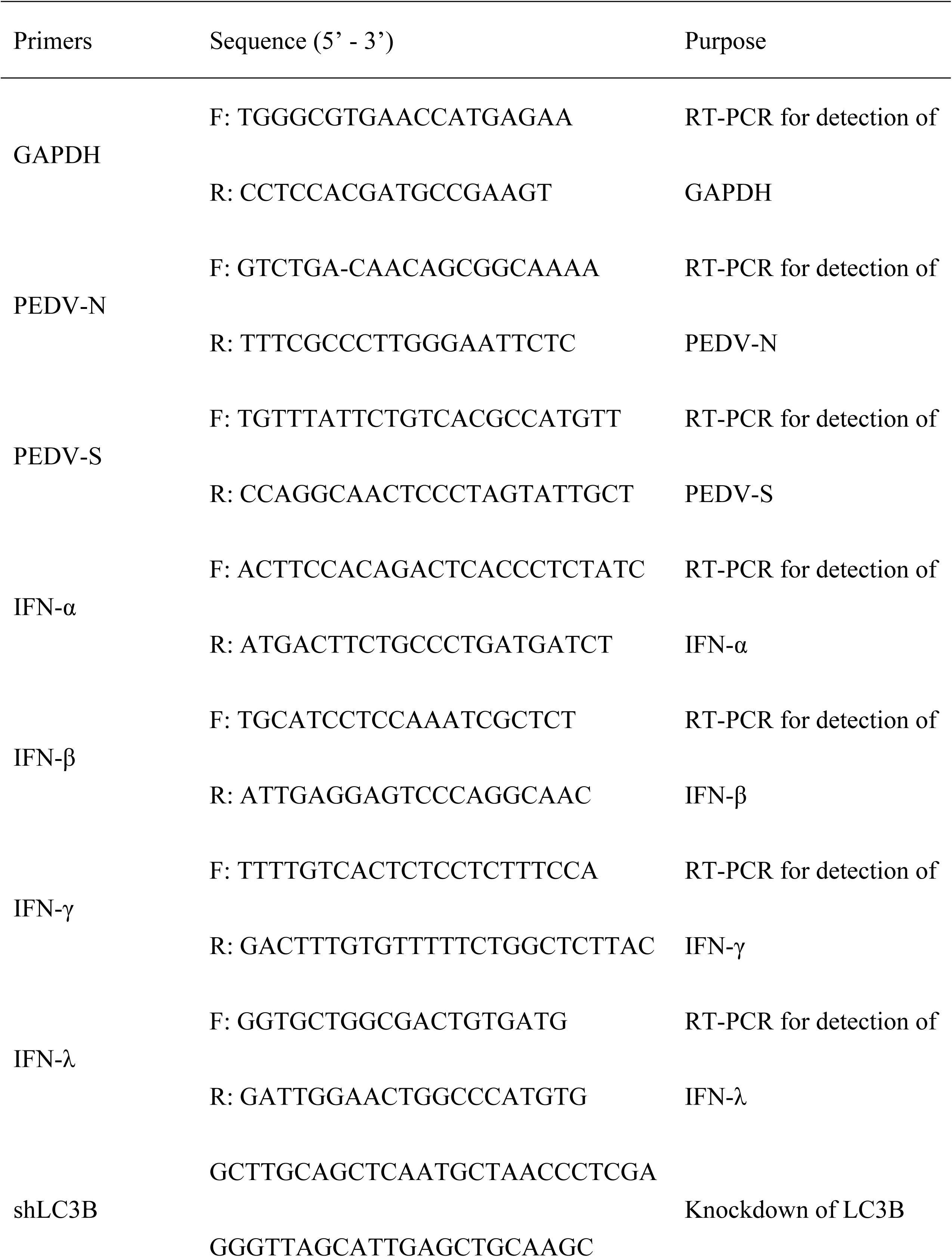
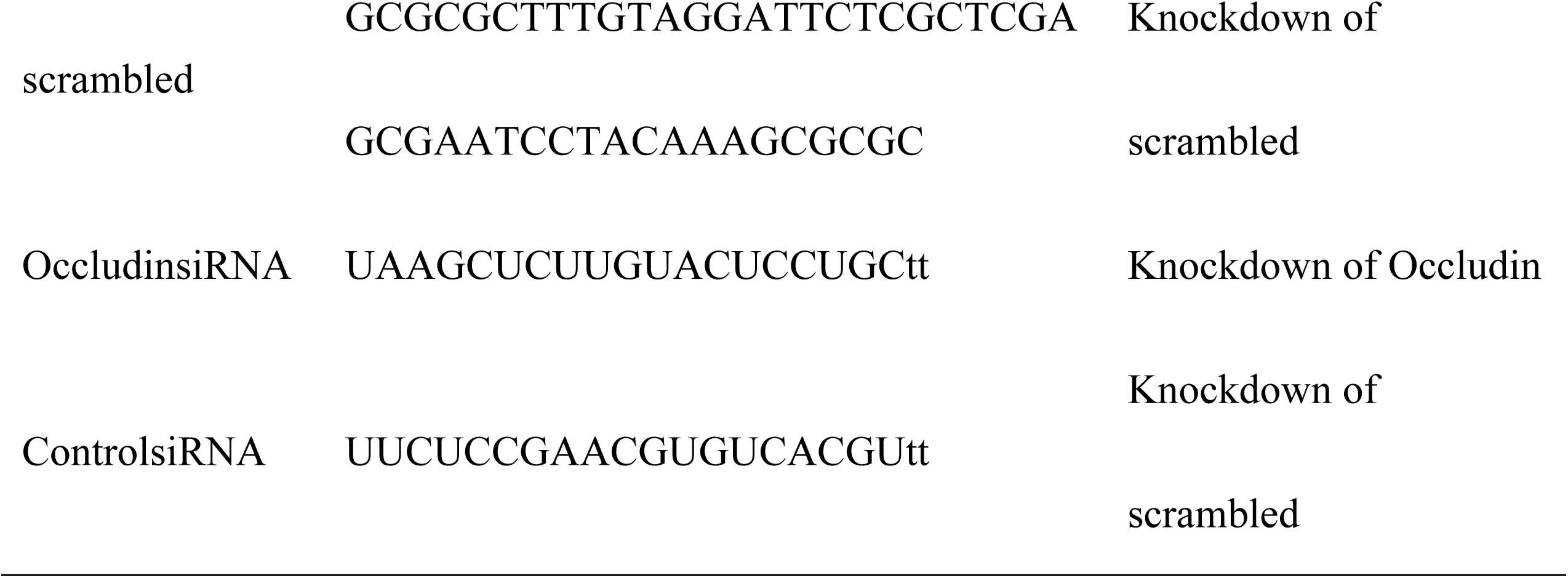
Primers used in this study.

### Immunoblotting analysis

The relative protein expression levels were measured by immunoblotting as previously described with minor modifications [51]. Briefly, equal amounts of protein obtained from the lysed cells were loaded onto 12% SDS-PAGE gels and transferred onto PVDF membranes (Millipore, USA). After blocking with 5% BSA for 2 h, the PVDF membranes were incubated at 4 °C overnight with primary antibodies, followed by a 1-h incubation with secondary antibodies at room temperature. The expected protein bands were detected using Image Quant LAS 4000 (GE Healthcare Life Sciences, USA). The relative abundance of the target protein (normalized to β-actin) was quantified by densitometric analysis using the Image Pro-Plus 6.0 software.

### Cell cultures

The porcine intestinal cell line IPEC-J2 cells were stored in our laboratory and grown in DMEM/F12 (1:1) medium supplemented with 10% fetal bovine Serum (Invitrogen, Carlsbad, USA), 1% insulin-transferrin-selenium (ITS), 5 ng/mL epidermal growth factor (EGF; Sigma, USA) and 1% antibiotics at 37 °C in a humidified atmosphere containing 5% CO_2_.

### Cell viability assay

Cell viability was monitored by 3-(4,5-dimethyl-2-thiazolyl)-2,5-diphenyl-2-H-tetrazolium bromide (MTT; Sigma, USA) assay as previously described [52]. Briefly, IPEC-J2 cells were cultured in 96-well plates at a density of 5 × 10^3^ cells/well with corresponding treatments. Then, each well was added with 15 μl of MTT (5 mg/ml) for another 4 h at 37 °C. The supernatants were discarded and incubated with 150 μl DMSO to dissolve the precipitate. Absorbance was measured at 490 nm with a reference wavelength of 595 nm. All tests were performed three times.

### LC3B*^−/−^* IPEC-J2 cell production by CRISPR/ Cas9 system

The small guide RNAs (sgRNAs) were designed using Breaking-Cas (http://bioinfogp.cnb.csic.es/tools/breakingcas/) online tool and synthesized (Invitrogen). The sgRNA was cloned pCas-Puro-U6 plasmid and the pCas-Puro-U6 plasmid Linear was obtained using the BbsI restriction enzyme (Thermo Fisher Scientific). The plasmids containing sgRNA were transfected into IPEC-J2 cells with GeneTran III (Biomiga) for 48 h, and then the transfected cells were selected using 5 μg/mL of puromycin. The selected cells were subjected to serial dilutions in 96-well plate to obtain a single cell colony. After 14 days of colony formation, each single colony was picked and expanded. Genomic DNA was extracted from individual clones and sequenced to confirm the specificity of targeting.

### Fluorescence Microscopy

Cells grown on coverslips were fixed with 4% paraformaldehyde for 20 min at 4 °C. After washing three times, cells were blocked with 1% BSA at room temperature and incubated with primary rabbit anti-occludin antibody and secondary FITC-conjugated goat anti-rabbit antibody (Invitrogen, USA), respectively. For the analysis of LC3B expression, cells grown on coverslips to 60-70% confluence were transfected with the pLVX-mRFP-EGFP-LC3B plasmid (provided by Prof. Qian Yang, Nanjing Agriculture University, Nanjing, China) using jetPRIME transfection reagent (Plolyplus-transfection, Illkirch, France) according to the manufacturer’s protocols. Nuclei were stained with DAPI (Blue, Beyotime Biotechnology, China). Fluorescence microscopy was performed using a Zeiss LSM710 confocal microscope (Zeiss, Oberkochen, Germany).

### RNA interference

Occludin-specific siRNA and control siRNA were designed and synthesized by Invitrogen (Thermo Fisher, USA). All primers are listed in Table 1. Cells were transfected with 100 nM occludin-specific or control siRNA duplexes by use of jetPRIME transfection reagent according to the manufacturer’s guidelines. Twenty-four hours after transfection, cells were washed with DMEM/F12 and cultured in DMEM/F12 with 4% FBS until further treatments.

### Quantification of virus titer

Viral titers were determined by 50% endpoint dilution (50% tissue culture infective dose [TCID50]) assays on IPEC-J2 cells as previously described [52]. Briefly, IPEC-J2 cells cultured in 96-well plates were inoculated with 10-fold dilutions of the harvested culture supernatants for indicated time. Microscope was used to detect the viral antigen according to the cell damage. Viral titers were expressed as TCID50/ml by using the Reed-Muench method.

### Flow Cytometry

Apoptotic cell death was measured by Annexin V/propidium iodide (PI) staining assay (BD Pharmingen, USA) using flow cytometry (FACS Calibur, BD Biosciences, USA) according to manufacturer’s instructions. In a word, the harvested cells were resuspended in 100 μl binding buffer followed by incubation with 10 μl Annexin V per test for 10 min, and 10 μl PI per test was added for 5 min. Cells were then suspended in 500 μl of binding buffer and immediately analyzed by FACS.

### Statistical Analysis

Statistical analyses were performed using Graph Pad Prism 7.0 by one-way analysis of variance (ANOVA), and the data were expressed as the means ± standard deviation (SD). *P* < 0.05 was regarded as significant.

## Acknowledgements

This study was financially supported by the National Key Research and Development Program (2017YFD0501001), the National Natural Science Foundation of China (31772811 and 31602123) and the Priority Academic Program Development of Jiangsu Higher Education Institutions (Jiangsu, China).

